# Nasal immunization with the C-terminal domain of BclA3 induced specific IgG production and attenuated disease symptoms in mice infected with *Clostridioides difficile* spores

**DOI:** 10.1101/2020.05.31.125435

**Authors:** Ana Raquel Maia, Rodrigo Reyes-Ramírez, Marjorie Pizarro-Guajardo, Anella Saggese, Ezio Ricca, Loredana Baccigalupi, Daniel Paredes-Sabja

**Affiliations:** Dipartimento di Biologia, Università di Napoli Federico II, Via Cinthia 26, 80126 Napoli, Italy; (A.R.M.); (A.S.); (E.R.); Microbiota-Host Interactions and Clostridia Research Group, Departamento de Ciencias Biológicas, Facultad de Ciencias de la Vida, Universidad Andrés Bello, Avenida Republica 330, 8370186 Santiago, Chile; (R.R.-R.); (M.P.-G.); Millennium Nucleus in the Biology of Intestinal Microbiota, Avenida Republica 330, 8370186 Santiago, Chile; Dipartimento di Medicina Molecolare e Biotecnologie Mediche, Università di Napoli Federico II, Via Pansini 5, 80131 Napoli, Italy

## Abstract

*Clostridioides difficile* is a Gram-positive, spore-forming bacterium that causes a severe intestinal infection. Spores of this pathogen enter in the human body through the oral route, interact with intestinal epithelial cells and persist in the gut. Once germinated, the vegetative cells colonize the intestine and produce toxins that enhance a strong immune response that perpetuate the disease. Therefore, spores are major players of the infection and ideal targets of new therapeutic treatments. In this context, spore surface proteins of *C. difficile*, are potential antigens for the development of vaccines targeting *C. difficile* spores. Here we report that the C-terminal domain of the spore surface protein BclA3, BclA3_CTD_, was identified as an antigenic epitope, over-produced in *Escherichia coli* and tested as an immunogen in mice. To increase antigen stability and efficiency, BclA3_CTD_ was also exposed on the surface of *B. subtilis* spores, a well-established mucosal vaccine delivery system. In the experimental conditions used in this study, free BclA3_CTD_ induced antibody production in mice and attenuated some CDI symptoms after a challenge with the pathogen, while the spore-displayed antigen resulted less effective. Although dose regimen and immunization route need to be optimized, our results suggest BclA3_CTD_ as a potentially effective antigen to develop a new vaccination strategy targeting *C. difficile* spores.

## Introduction

*Clostridioides difficile* is a Gram-positive, spore-forming and obligate anaerobe gastrointestinal bacterium, responsible for the most common nosocomial infection in industrialized countries [1]. In recent years the incidence and severity of *C. difficile* infections (CDI) has increased worldwide due to the emergence of antibiotic-resistant and hyper-virulent strains. In addition, about 20% of the infected people develop a second CDI episode within 2 months and in the case of more than two episodes the frequency of further recurrences increases dramatically up to 60% [2-4]. Nowadays, CDI is a major concern and an economic burden. Recent data indicated that CDI is more common than methicillin-resistant *Staphylococcus aureus* infections [5] and estimated a cost per CDI episode ranging from 5.000 to 12.000 euros in the European Union [6] and approximately 21.000 US dollars in the United States [7].

CDI is mainly transmitted by *C. difficile* spores through the fecal-oral route. Ingested spores survive the transit through the stomach, interact with intestinal epithelial cells and persist in the host gut. When gut conditions are favorable, i.e. when the number of other members of the gut microbiota is severely reduced, *C. difficile* spores germinate and massively colonize the gut. Growing cells of *C. difficile* then produce virulence factors that induce a strong immune response and the symptoms associated to *C. difficile* infections. Being infection vehicles, mediators of the initial interaction with intestinal cells and responsible of the persistence of the pathogen in the animal gut, *C. difficile* spores are key players of CDI and ideal targets of anti-*C. difficile* therapeutic treatments.

It is known that the hydrophobicity of *C. difficile* spores, due to the proteinaceous exosporium, contributes to adhesion to hospital surfaces and to intestinal epithelial cells (IECs) [8, 9] and members of the BclA family of collagen-like glycoproteins are abundantly present in the *C. difficile* exosporium. In the hyper-virulent strain R20291 of *C. difficile* the BclA proteins showed 56% similarity with the BclA protein of *Bacillus anthracis* [10], known to be highly immunogenic and to act as spore surface ligand for α2β1 integrin present in IECs driving spore entry into the epithelial barrier [11]. Further investigation needs to be done in order to fully understand the role of BclA proteins in *C. difficile* spores however, strong evidence suggest that these proteins are involved in the formation of the hair-like projections in most *C. difficile* strains such as the hyper-virulent strain R20291 [12, 13] and their morphology and localization proposes them as potential antigens [14]. While the antigenicity of BclA1 and BclA2 has been recently tested [15-18], BclA3 has not been directly evaluated to date.

BclA3 is composed by an N-terminal domain, possibly oriented to the inside, a collagen-like domain formed by GXX repeats, which is highly glycosylated [10], and a C-terminal domain that is presumably faced outwards of the exosporium [17]. Mutant *C. difficile* spores lacking BclA3 have shown a reduced colonization efficiency in a mice infection model [16] and a recent study has shown that several glycosylated peptides from BclA3 were able to induce humoral immune response in mice [19]. These results suggest that BclA3 glycoprotein could be an interesting antigen in a *C. difficile* spore-based vaccination strategy.

In this study, the BclA3 amino acid sequence was analysed *in silico* and the C-terminal domain, BclA3_CTD_, was identified as a potential epitope. BclA3_CTD_ antigenicity was then tested *in vivo* in a murine model as a free protein or displayed on *Bacillus subtilis* spores, a well-established antigen delivery system [20], proved to efficiently interact with antigen presenting cells (APCs) leading to the induction of humoral, local and cellular responses [21, 22]. Mice immunized with recombinant spores or with the pure antigen were able to produce BclA3_CTD_–specific IgG. The immunization with pure BclA3_CTD_ also impaired weight loss after a challenge with *C. difficile* spores and induced a decrease on *C. difficile* spore load in feces one day after the infection.

## Results

### *In silico* analysis of BclA3 and construction of the recombinant strain expressing the chimera protein CotBΔ-BclA3_CTD_

To predict the most immunogenic domain of BclA3 we used the Immune Epitope Database (IEDB) to analyse BclA3 amino acid sequence and predict continuous linear B and T cells MHC-I and MHC-II epitopes. As shown in Figure 1A, the C-terminal 170 amino acid residues of BclA3 showed the highest B cell antigenicity score. The prediction of T cell epitopes also suggests this part of the protein as the most immunogenic (data not shown). Based on that and on a previous report suggesting that the C-terminus of BclA3 is faced outwards of the exosporium of *C. difficile* spores [17], we selected the C-terminal part of BclA3 (BclA3_CTD_) (Figure 1B) as a putative antigen to be tested *in vivo* as a free protein and for display on the surface of *B. subtilis* spores. Hence, His-tagged BclA3_CTD_ was overexpressed in *Escherichia coli* BL21(DE3) and purified by affinity chromatography with Ni-sepharose columns as described in the Methods section. Upon purification, the protein was loaded on an SDS-Page gel (Figure 1C) next to pure BclA2_CTD_, an exosporium protein already purified by our group [18]. Whilst pure BclA2_CTD_ migrates on SDS-Page gel as a single band of approximately 14 kDa, as expected, pure BclA3_CTD_ consistently migrates in two bands, one with about 20 kDa and other with approximately 60 kDa that match with the predicted sizes of a monomer and trimers, respectively, which may indicate protein aggregation.

**Figure 1.**
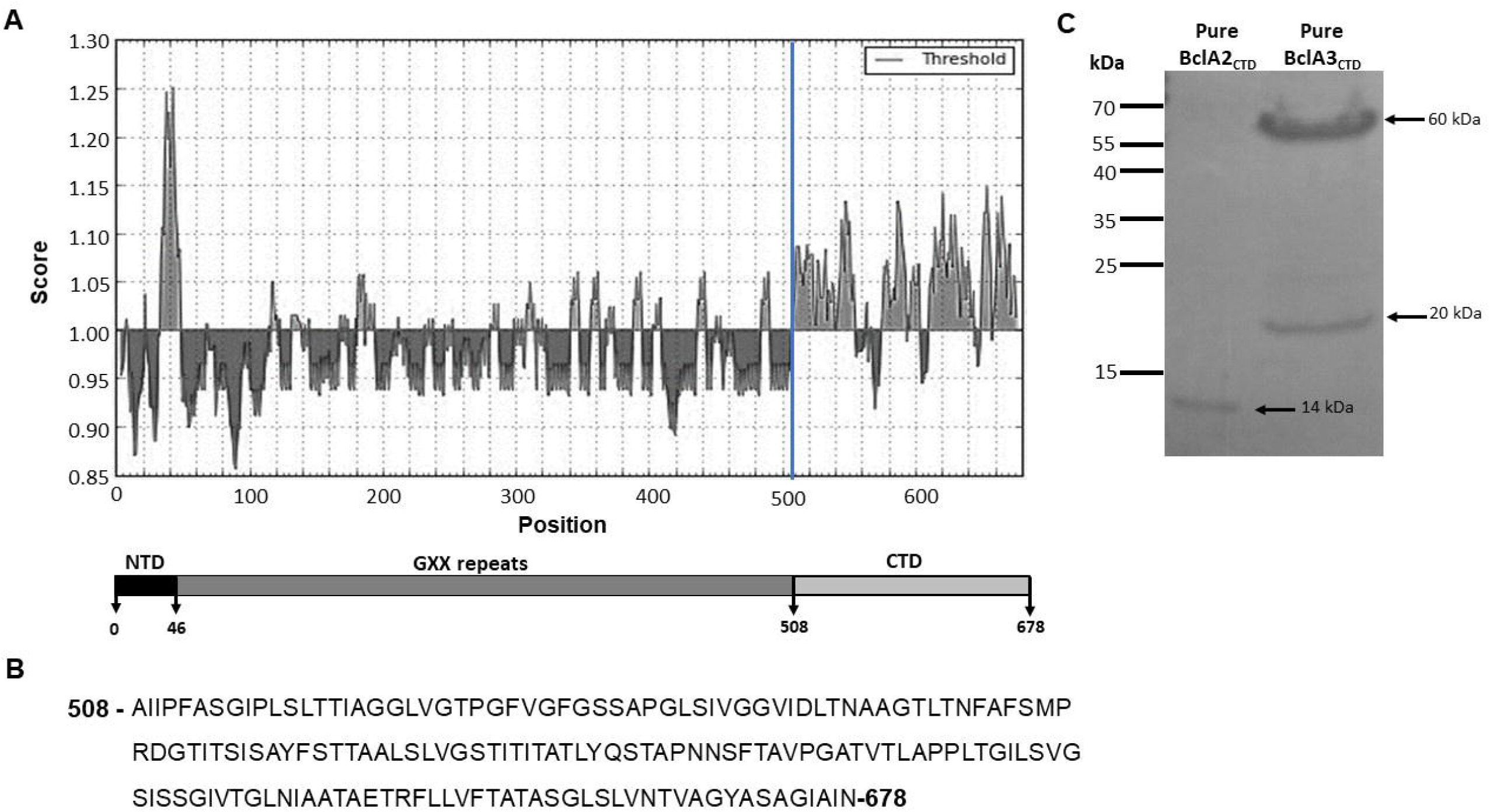
Analysis of the glycoprotein BclA3 from *C. difficile* R20291. **(A)** The C-terminal domain (CTD) of BclA3 (last 170 amino acid residues) shows higher B cell epitope propensity score compared with the rest of the protein (Kolaskar & Tongaonkar Antigenicity Method from Immune epitope database). The X- and Y-axes represent the sequence position and antigenic propensity score, respectively. The threshold value was generated by default by Immune epitope database (http://tools.iedb.org/bcell/). The regions above the threshold are antigenic. **(B)** The correspondent aminoacidic sequence of BclA3_CTD_ is shown. **(C)** Coomassie stained SDS-Page gel. 2 and 5 µg of purified C-terminal domains of the exosporium proteins BclA2 and BclA3 were loaded, respectively. BclA2CTD migrates as 14 kDa band while BclA3_CTD_ migrates as 2 bands of approximately 20 and 60 kDa.

DNA coding for the last 170 amino acid residues of BclA3 (BclA3_CTD_) was used not only to over-express and purify the protein fragment but also to construct a gene fusion with DNA coding for the *B. subtilis* spore surface protein CotB, a coat protein already used to anchor heterologous antigens [23-27]. In particular, we used a truncated version of CotB, CotBΔ, by deleting 105 C-terminal amino acids, thus removing a region with repeated sequences avoiding potential structural instability of the genetic constructs [23]. As previously reported [23], the gene fusion was cloned into an integrative vector adjacent to a chloramphenicol-resistance gene cassette (Cm^R^) and used to transform competent cells of the *B. subtilis* strain PY79 [28]. Chloramphenicol-resistant clones were the result of a double cross-over integration event, schematically indicated in Figure 2A. Chloramphenicol-resistant clones were tested for the site of chromosomal integration by PCR (not shown) and clone AZ703 was selected for further analysis.

**Figure 2.**
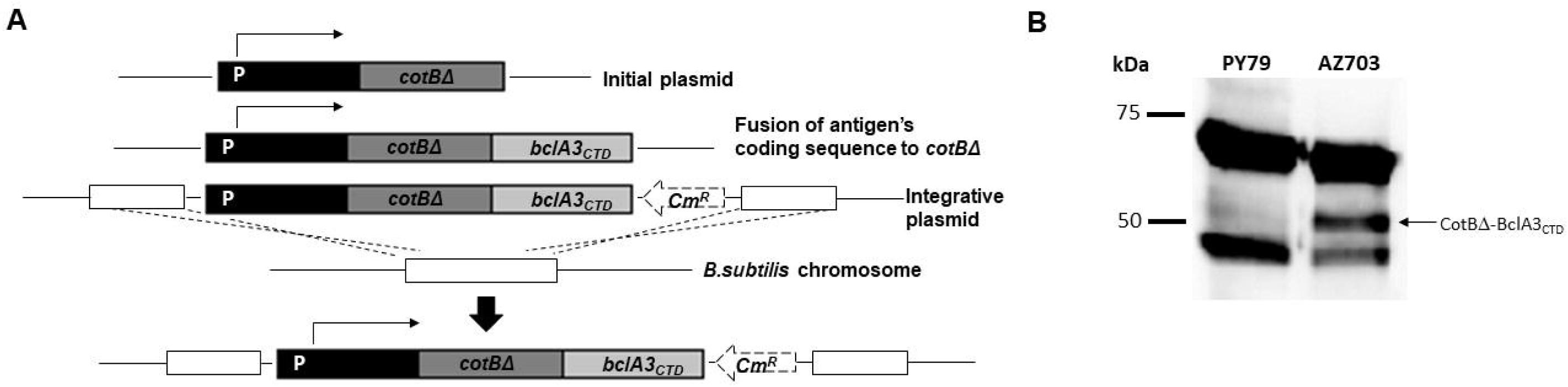
Construction of *B. subtilis* recombinant strain AZ703. **(A)** Schematic representation of the strategy for the integration of the gene fusion *cotBΔ::bcla3_CTD_* in the chromosomal DNA of *B. subtilis* PY79. The western-blot analysis of proteins extracted from spores of *B. subtilis* laboratory strain PY79 (lane 1 in **B**) and from the recombinant strain AZ703 (lane 2 in **B**) show bands of 66 and 46 kDa, which correspond to the endogenous CotB protein. The lane of recombinant strain AZ703 also displays a band of about 50 kDa which corresponds to the chimera protein CotBΔ-BclA3_CTD_. 5×10^8^ of spores were resuspended in 100 µL of loading buffer and 20 µg of protein extract was loaded in SDS-Page gel. The immunoreaction was performed with anti-CotB antibodies and anti-rabbit secondary antibody conjugated with horseradish peroxidase.

Purified spores of strain AZ703 were used to extract surface proteins by the SDS-DTT procedure [29] and extracted proteins were analyzed by Western-blot with anti-CotB antibody. CotB has a deduced molecular mass of 46 kDa but it is known to migrate on SDS-page in two forms: a predominant form of 66 kDa and a minor form of 46 kDa [30]. Both forms were extracted from PY79 and AZ703 spores with only the latter also showing an additional protein, slightly bigger than 50 kDa (Figure 2B). The additional protein was recognized by the anti-CotB antibody and conformed well with the expected size for the fusion protein, since the truncated form of CotB and the BclA3 fragment have predicted sizes of 36 kDa [23] and 20 kDa (Figure 1C), respectively.

### Mice intranasal immunization

A mucosal immunization experiment was performed in a murine model to test the efficiency of BclA3_CTD_ as an antigen, both as a free protein and upon display on *B. subtilis* spores. Mice were divided into four experimental groups and nasally immunized 3 times either with PBS pH 7 (n=11), 2×10^9^ spores of *B. subtilis* PY79 (n=11) (Sp), 2×10^9^ spores of AZ703 (n=11), or 4 μg of purified BclA3_CTD_ (n=11). The animal serum was collected one day before the first (pre-immunization day, PI), second (day 13) and third (day 27) immunizations as well as two days before *C. difficile* infection (day 42). One day before the infection with 5×10^7^ spores of the *C. difficile* strain R20291, mice were treated with Clindamycin as previously reported [31] and schematically shown in Figure 3.

**Figure 3.**
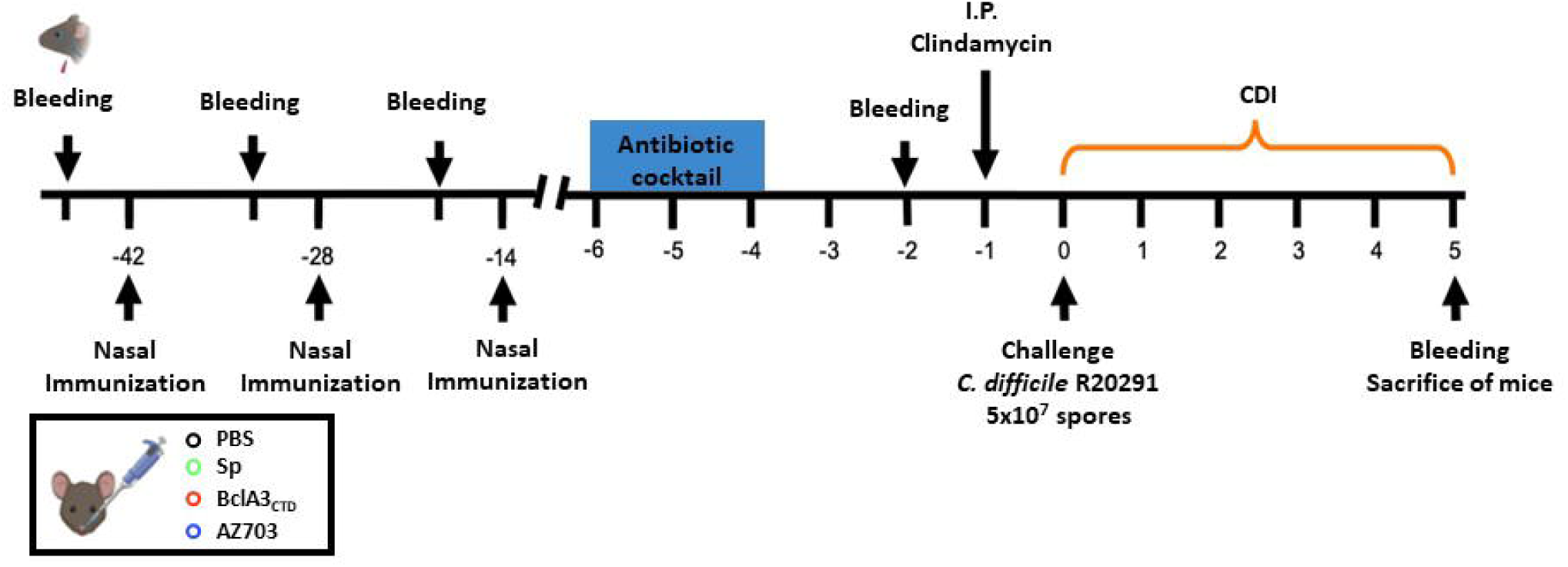
Overview of the experimental design schematics for the prevention of *C. difficile* infection in a murine model. C57BL/6 mice were nasally immunized 3 times (42, 28 and 14 days before challenge with *C. difficile* R20291 spores) with PBS, spores of *B. subtilis* PY79, purified BclA3_CTD_ or spores of *B. subtilis* displaying the chimera protein CotBΔ-BclA3_CTD_ (AZ703). Prior to the infection with *C. difficile* R20291, the animals were submitted to an antibiotic cocktail (days 4-6 before challenge) and clindamycin administration (1 day before challenge). On day 0 mice were infected with 5×10^7^ of *C. difficile* R20291 spores and were monitored from day 0 to day 5 for CDI symptoms. Serum was collected one day before each immunization, two days before *C. difficile* infection and on the day of sacrifice.

BclA3_CTD_ immunogenicity was measured by ELISA analysing the presence of anti-BclA3_CTD_ IgG in animal serum throughout the experiment. As shown in Figure 4A mice immunized with pure BclA3_CTD_ produced BclA3_CTD_-specific IgG upon nasal administration even on the 13^rd^ day (immunized only once). After the second and third immunizations (d27 and d41, respectively) the increase was significant (P < 0.0001 compared to Pre-immune serum and with d13). The recombinant spores AZ703 also induced significant levels of anti-BclA3_CTD_ IgG after the third immunization (P = 0.0380 compared to Pre-immune serum). Neither PBS nor *B. subtilis* PY79 spores were able to raise an anti-BclA3_CTD_ IgG response in mice, as expected.

**Figure 4.**
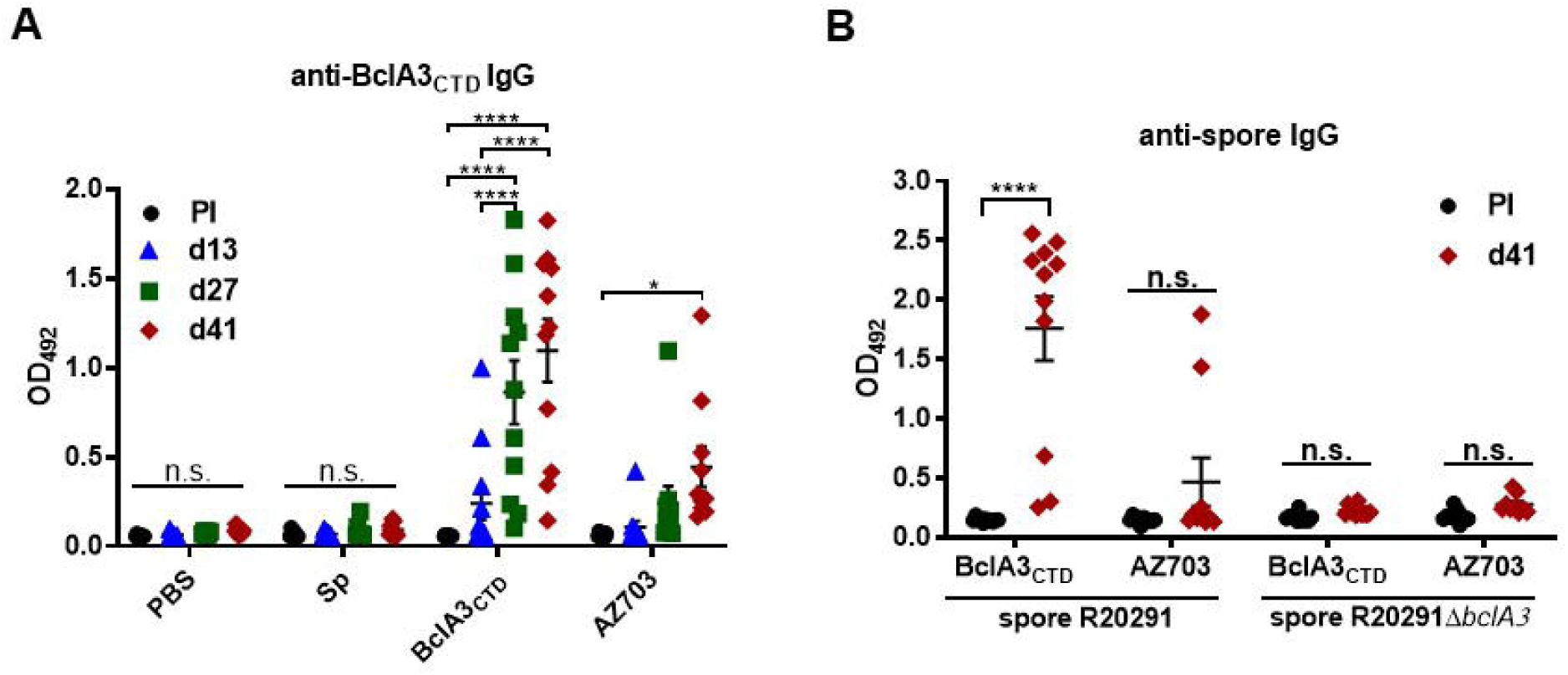
Immunogenicity of pure BclA3_CTD_ protein and AZ703 spores in mice nasal immunization. **(A)** IgG anti-BclA3_CTD_ was measured in mice serum on days 0 (Pre-Immune serum, PI), 13 (d13), 27 (d27) and 41 (d41). **(B)** Anti-*C. difficile* R20291 spores or anti-*C. difficile* R20291Δ*bclA3* spores IgG levels were measured in the serum of mice on day 0 and 41. Serum collection on day 0, 13 and 27 was made one day before each immunization and on day 41 two days before infection. Results were determined by ELISA and are reported as optical density (OD) units at 492nm. The geometric mean plus standard error of the mean for each cohort are shown. Comparisons between days in the same group were obtained using Two-way ANOVA Turkey’s multiple comparison test and statistical significance (P < 0.05) is indicated by asterisks. n.s means no significance.

The immune reactivity of the serums collected 2 days before *C. difficile* R20291 infection when incubated either with spores of the hyper-virulent strain R20291 or with the isogenic *bclA3* mutant strain (R20291Δ*bclA3*) was also tested. As shown in Figure 4B, mice immunized with pure BclA3_CTD_ produced IgG able to recognize R20291 spores (P < 0.0001 in comparison to the Pre-immune serum) but not R20291Δ*bclA3* spores indicating the specificity of the response. As expected by the low immune response induced by spore-displayed BclA3 (Figure 4A), only two out of eleven mice immunized with spore-displayed BclA3_CTD_ produced IgG able to specifically recognize R20291.

In conclusion, results of Figure 4 show that BclA3_CTD_ is an antigen able to induce the production of BclA3_CTD_-specific IgG in a murine model and serum of animals immunized with the pure antigen were also able to recognize spores of the hypervirulent strain R20291 of *C. difficile*. When displayed on *B. subtilis* spores BclA3_CTD_ is still able to induce the production of BclA3-specific IgG, even if at a lower level.

### Effect of nasal-BclA3_CTD_ immunization against *C. difficile* R20291 infection

Figure 5A shows that intranasal immunization with purified BclA3_CTD_ prevented weight loss after the challenge with 5×10^7^ spores of the R20291 strain of *C. difficile*. In particular, on days 1 and 2 post-infection a statistically significant difference (P = 0.0177 and P = 0.0099, respectively) was observed between mice immunized with pure BclA3_CTD_ and those immunized with wild-type spores of *B. subtilis* (Figure 5A). No statistically significant differences were observed concerning the appearance of diarrhea caused by the challenge with R20291 spores, indicating that the nasal immunization with pure BclA3_CTD_ or with the recombinant strain displaying CotBΔ-BclA3_CTD_ did not halt all CDI symptoms (Figure 5B). Plus, the severity of diarrhrea, associated with a high score (0 meaning normal stool and 3 liquid stool), did not vary significantly between groups in the same day (Figure 5C).

**Figure 5.**
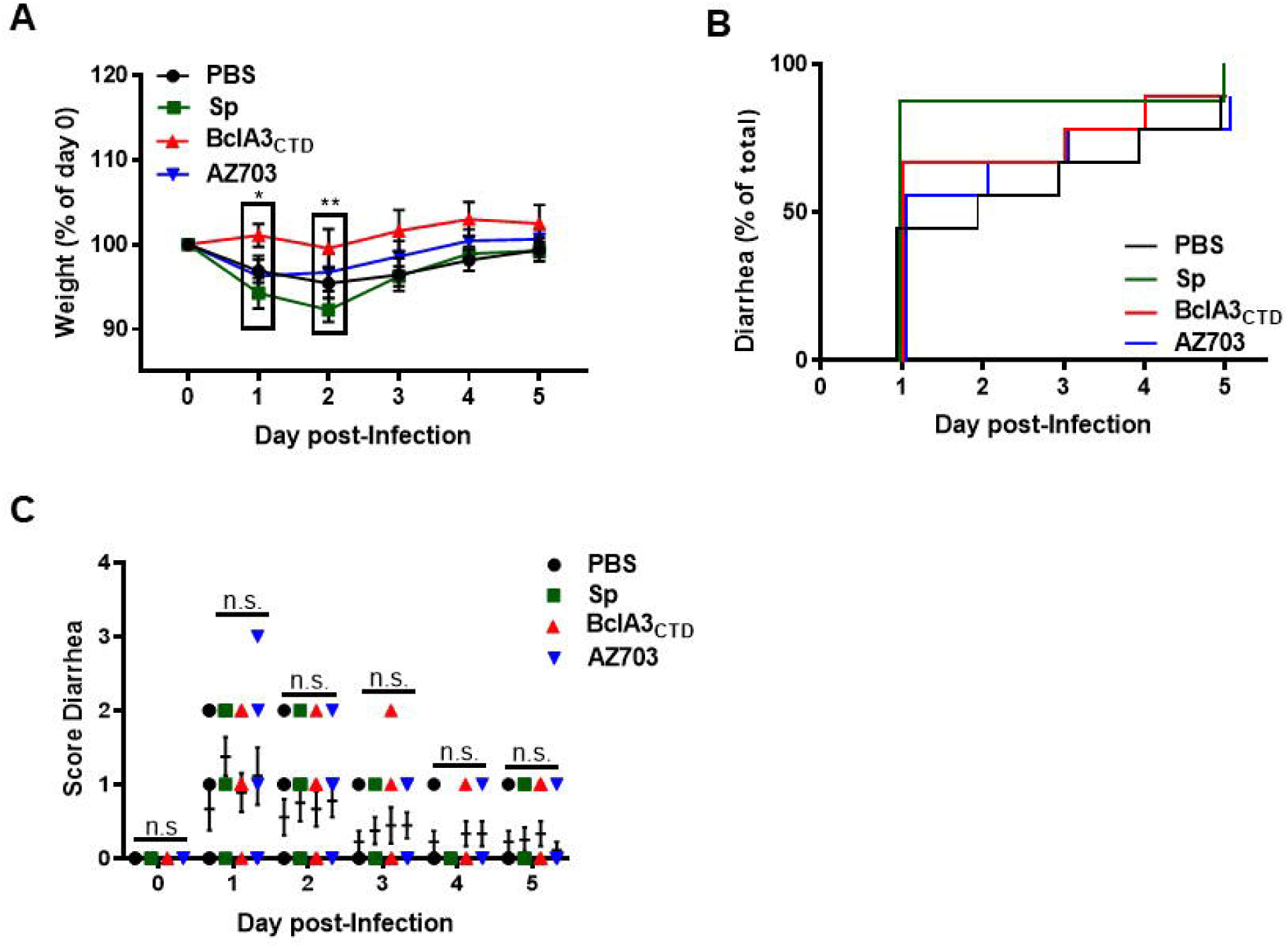
Protective effect of BclA3_CTD_ and AZ703 intranasally administered against CDI in murine model. C57BL/6 mice were nasally immunized with PBS, *B. subtilis* PY79 spores, purified BclA3_CTD_ or AZ703 spores and challenged with *C. difficile* R20291 spores. Mice were monitored in the following 5 days after infection for **(A)** Weight loss presented as the relative % of the weight to the day of infection (day 0 or D0); **(B)** Time of occurrence of diarrhea, presented as the relative % of diarrhea in a group to the total mice and **(C)** Score of diarrhea per day. Error bars are standard error of the mean. Two-way ANOVA Tukey’s multiple comparisons test (A and C); Log-rank (Mantel-Cox) test (B). Statistical significance (P < 0.05) is indicated by asterisks. n.s means no significance.

In order to evaluate if the immunization influenced *C. difficile* sporulation inside the host and spore clearance, we compared the spore levels present in stools within 5 days post-infection. We observed a statistically significant lower spore load in the feces of mice immunized with BclA3_CTD_ one day post-infection (P < 0.0001 with respect to mice immunized with PBS or PY79 spores and P = 0.0002 with mice immunized with AZ703 spores) (Figure 6A) suggesting that animals immunized with the pure antigen were able to quickly eliminate *C. difficile* spores. However, from day 2 to day 5 all other groups of mice were similarly able to eliminate *C. difficile* spores (Figure 6A). No differences were observed in the *C. difficile* spore load in the ileum, proximal, middle or distal colon tissue (not shown). Finally, we measured toxin levels in mice feces, a sign of *C. difficile* colonization inside the cecum. As shown in Figure 6B, no statistically significant differences were observed suggesting that the immunization strategies could not prevent colonization and cytotoxicity in the cecum.

**Figure 6.**
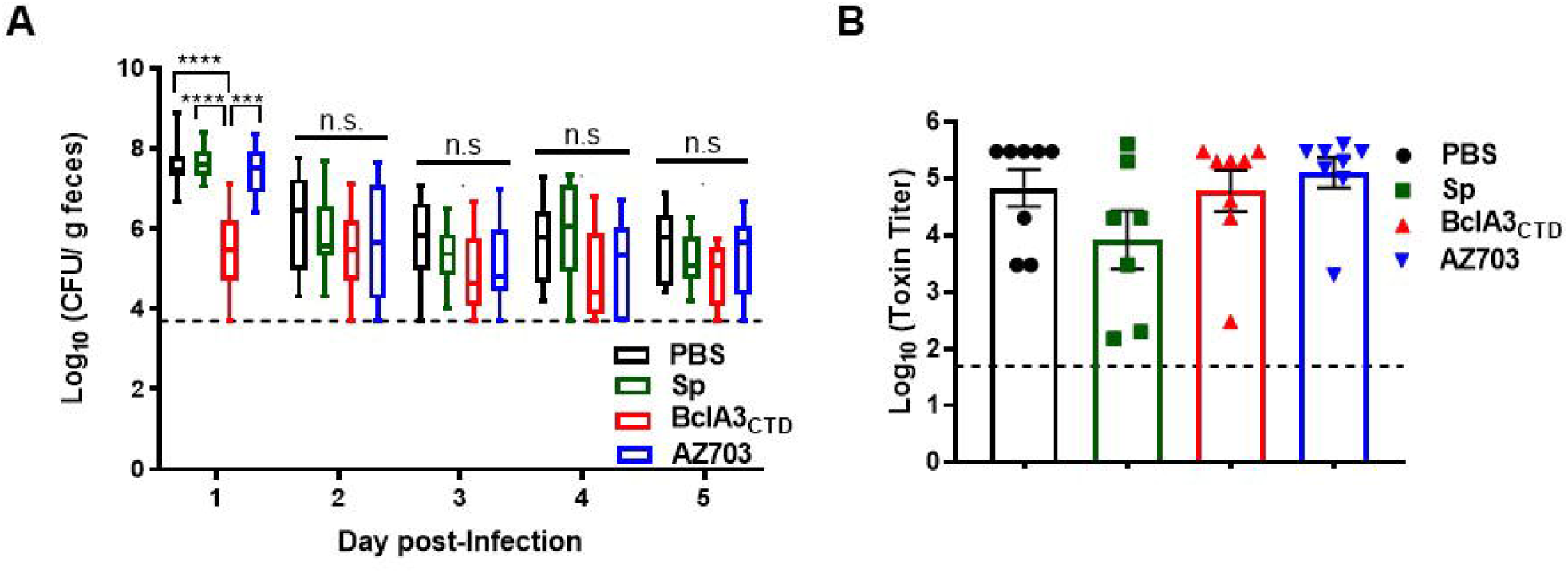
Analysis of spore load in feces and cecal toxin titers. **(A)** The load of *C. difficile* spores in the feces was evaluated on the following 5 days after infection as log10 CFU/g of feces. **(B)** The cecum content toxicity was measured and represented as log_10_ toxin titer. Two-way ANOVA Tukey’s multiple comparisons test in A and Mann-Whitney non-parametric test in B were used; Statistical differences (P < 0.05) are indicated by asterisks. The bars are the geometric mean ± standard error of the mean. n.s means no significance.

## Discussion

Current treatment options for CDI rely on the use of antibiotics, fecal microbiota transplantation, probiotic administration or monoclonal antibodies against *C. difficile* toxins [32]. The spore, form in which *C. difficile* persists inside the host, possibly modulating its immune system, is still not considered as a main target for therapies. We have recently demonstrated that the C-terminal domain of BclA2 elicits an elevated humoral response [18]. Here, we have identified as a potential antigen to induce an anti-spore immune response, the C-terminal domain of the exosporium protein BclA3 (BclA3_CTD_) of the hyper-virulent strain R20291 of *C. difficile*.

Here we show that mice nasally immunized with pure BclA3_CTD_, even only two times, were able to produce BclA3_CTD_-specific IgG, indicating that the antigen was able to elicit a specific humoral immune response and therefore could be exploited for a vaccination strategy against *C. difficile*. Moreover, upon challenge with *C. difficile* spores, mice immunized with pure BclA3_CTD_ not only maintained their weight but also had a reduction in *C. difficile* spore load in feces even one day after the infection. Considering an induction of a robust specific systemic immune response, as suggested by the high titers of IgG anti-BclA3_CTD_, it is tempting to assume that animals immunized with the pure peptide have activated the complement signaling cascade leading to the opsonization and neutralization of *C. difficile* spores, phagocytic elimination and therefore experienced spore clearance earlier than the other groups of animals [33, 34]. However, it is noteworthy that the immunization with pure BclA3_CTD_ was not able to halt neither the occurrence and intensity of diarrhea nor to reduce the spore load in colonic tissues or to avoid *C. difficile* spore colonization in mice cecum. These results may indicate that the systemic immune response was not strong enough to prompt local immunity and a complete protective immune response. The change of administration route and/or optimization of antigen-dose could possibly overcome these limitations. Plus, the measurement of mucosal IgA will be addressed in the following experiments for a better understanding of local immune responsiveness.

We have also immunized mice with *B. subtilis* spores recombinantly engineered to display BclA3_CTD_ anchored to a highly abundant coat protein, CotB. Due to its safety and robustness *B. subtilis* spores have been widely used as mucosal vaccine vehicles [35-37]. *B. subtilis* spores administered by the nasal route have been shown able to expose the antigen to mucosal-associated lymphoid tissue (MALT) and therefore prompt a strong immune response [38-40]. Plus, it is shown that antigens are more protected from enzymatic lysis and have increased stability when displayed into *B. subtilis* spore surface than free antigen [21, 41]. Nevertheless, our work only showed a slight increase of specific anti-BclA3_CTD_ IgG in mice immunized with recombinant spores and only after three immunizations. Consequently, it is not surprising that we have not observed and impairment in CDI symptoms. Preliminary work developed by our group (data not shown) suggests a low yield of the chimera protein CotBΔ-BclA3_CTD_ on the surface of *B. subtilis* spores and, as a consequence, a low dose of antigen administered to mice. Further experiments to go deeper in this subject are still in progress. Yet, we consider promising the weak immune response observed in the experiments reported here and plan to explore the possibility of using higher doses of spore-delivered antigen and of using the oral route of immunization to increase the efficacy of spore-displayed BclA3. In this context, it is known that recombinant spores of *B. subtilis* administered by gavage germinate in the stomach and small intestine and due to low oxygen present in the large bowel environment they re-sporulate displaying the heterologous antigen again [42-44].

In conclusion, the induction of humoral immune response and the partial protective effects observed in animals nasally immunized with the purified protein BclA3_CTD_, clearly indicate that BclA3_CTD_ is a promising antigen to be tested in future *in vivo* trials.

## Materials and Methods

### Bacterial strains and spore purification

*E. coli* strains DH5α and BL21 (DE3) (Invitrogen) were used for cloning and BclA3_CTD_ overexpression, respectively. *B. subtilis* PY79 [28] was used as a as a parental strain of AZ703. The hyper-virulent strain R20291 of *C. difficile* was used for mice infection. To test BclA3_CTD_-specific immunogenicity an R20291Δ*bclA3* knock-out mutant was used (Paredes-Sabja, unpublished work).

Sporulation of *B. subtilis* PY79 and AZ703 was induced by the exhaustion method [45]. Briefly, after 35 hours of growth in Difco Sporulation (DS) medium at 37°C with vigorous shaking, spores were collected, washed and purified. The purification was performed using KCl 1 M, lysozyme 10 mM, NaCl 1 M, SDS 0,05% and several washes with water.

*C. difficile* spores were purified as described elsewhere [46]. Spore suspensions were prepared by plating a 1:100 dilution of an overnight culture onto a 70:30 medium (63 g Bacto peptone (BD Difco), 3.5 g proteose peptone (BD Difco), 0.7 g ammonium sulfate (NH_4_)_2_SO_4_, 1.06 g Tris base, 11.1 g brain heart infusion extract (BD Difco) and 1.5 g yeast extract (BD Difco) for 1 L) and incubating it for 5 days at 37°C under anaerobic conditions [47]. After incubation, plates were removed from the chamber and the surface was scraped up with ice-cold sterile water. Next, the spores were washed five times gently with ice-cold sterile water in micro centrifuge at 14 000 rpm for 5 minutes. Spores were loaded onto a 45% Nycodenz solution, centrifuged (14 000 rpm, 40 minutes). After centrifugation, the spore pellet was washed five times (14 000 rpm, 5 minutes) with ice-cold sterile water to remove Nycodenz remnants.

The spores were counted in Neubauer chamber and volume adjust at 5×10^9^ spores per mL. Spore suspensions were purified until they were >99% free of vegetative cells, sporulating cells and cell debris as determined by phase-contrast microscopy.

### BclA3_CTD_ over-production and purification

The chromosomal DNA of *C. difficile* R20291 was used for the amplification of *bcla3* C-terminal domain (CTD) (513 bp) with the oligonucleotides BclA3_CTDsense_ (ggtaccggatccGCAATAATACCTTTTGCATCAGG, in lower case is the recognition site for KpnI, NcoI and BamHI restriction enzymes) and BclA3_CTDanti_ (tctagactgcagCTAATTTATTGCAATTCCTGCAC in lower case is the recognition site for XbaI and PstI restriction enzymes) to prime the reaction. The coding sequence of BclA3_CTD_ was cloned in the plasmid pGEMT-easy (Promega) and posteriorly cleaved with BamHI/PstI restriction enzymes and inserted in-frame to an N-terminal polyhistidine tag in the expression vector pRSETA (Invitrogen), previously digested with the same enzymes. Upon transformation of *E.coli* BL21(DE3) with pRSETA::*bcla3*_*CTD*_, the strain was incubated in ampicillin-supplemented (50 µg/mL) TY medium. Once reached an optical density of 0.7 at 600 nm the culture was added to an autoinduction medium (T7 promoter induction by lactose) and incubated for 16 hours at 37°C with shaking. The six-His-tagged BclA3_CTD_ protein was purified under native conditions using the His-Trap column as recommended by the manufacturer (GE Healthcare Life Science). The purified protein was desalted and concentrated with the Centricon cutoff 10 kDa (Merck Millipore). The purity of the protein was analyzed by SDS-page and Western-blot using Anti-His antibodies.

### Construction of the recombinant strain AZ703

DNA coding for BclA3_CTD_ and for the N-terminal 275 amino acids of CotB were PCR amplified using the *C. difficile* R20291 and *B. subtilis* PY79 chromosome as template, respectively. To prime *cotBΔ* the oligonucleotides B1 (acatgcatgcACGGATTAGGCCGTTTGTCC in lower case there is the recognition site for SphI restriction enzyme) and B3 (gaaagatctGGATGATTGATCATCTGAAG in lower case there is the recognition site for BglII restriction enzyme) were used. The obtained amplification products were cloned in pGEMT-easy (yielding pGEMT-easy::*bclA3*_*CTD*_ and pGEMT-easy::*cotBΔ*). The *bclA3*_*CTD*_ gene was digested from pGEMT-easy::*bclA3*_*CTD*_ with BamHI/PstI restriction enzymes and cloned in-frame to the 3′ end of the *cotBΔ* gene carried by plasmid pGEMT-easy::*cotBΔ* previously diggested with BglII/PstI, yielding plasmid pGEMT-easy::*cotBΔ*::*bclA3*_*CTD*_. The fusion *cotBΔ::bclA3*_*CTD*_ gene was digested with the restriction enzymes SphI/SalI and ligated to the previously digested integrative plasmid pDG364 which contains a coding region for chloramphenicol resistance. Competent *B. subtilis* PY79 cells were transformed with the previously linearized integrative vector with NdeI and plated in medium with chloramphenicol. The antibiotic-resistant clones were the result of double-crossover recombination with *amyE* gene on the *B. subtilis* chromosome. The chromosomal DNA was extracted from the positive clones and tested by PCR. Sporulation of PY79 and recombinant strain (AZ703) was induced by nutrient exhaustion in DS medium. After 35 hours incubation at 37°C, spores were collected, washed and purified as described before. The coat proteins from 5×10^8^ spores of PY79 and AZ703 were extracted by SDS-DTT treatment. To verify the expression of the chimera protein CotBΔ-BclA3_CTD_, extracted proteins were analyzed by western-blot using anti-CotB antibodies.

### Western-blot analysis

The coat proteins from 5×10^8^ spores of *B. subtilis* PY79 and AZ703 were extracted by SDS-DTT treatment [29] and quantified by Bradford assay (BioRad). 20 µg of protein extract or 500 ng of pure BclA3_CTD_ with protein sample buffer 2x [50] were incubated at 100°C for 7 minutes and loaded onto a 12% SDS-Page gel. Proteins were then electro-transferred to nitrocellulose filters (Amersham Pharmacia Biotech) and used for Western-blot analysis by standard procedures. To identify the recombinant protein CotBΔ-BclA3_CTD_ it was used anti-CotB 1:7000 as primary antibody and anti-rabbit secondary antibody conjugated with horseradish peroxidase 1:7000. To identify pure BclA3_CTD_ it was used the antibody anti-His 1:7000.

### Animals

Mice 8-12 weeks old C57BL/6 (male or female) were obtained from a breeding colony at Facultad de Ciencias Biologicas Universidad Andres Bello (Santiago, Chile), established using animals purchased from Jackson Laboratories. Water, bedding and cages were previously autoclaved and mice had a 12-hour cycle of light and darkness. All experimental protocols were conducted in strict accordance with and under the formal approval of the Biologicals Sciences Faculty of Universidad Andrés Bello.

### Immunization Regimen in mice

Mice were randomly assigned to four experimental groups (11 animals each group) according to the type of immunization received. Mice were intranasally immunized on days 0, 14 and 28 with 20 μL (10 µl per nostril) of PBS pH 7, 2×10^9^ spores of *B. subtilis* PY79, 2×10^9^ spores of AZ703 or 4 μg of purified BclA3_CTD_. The day before each immunization, two days before infection and on the day of the sacrifice blood was collected.

### Animal infection model

Prior to infection, mice were pre-treated with antibiotic cocktail of kanamycin (40 mg/kg body weight; Sigma-Aldrich, U.S.A.), gentamicin (3.5 mg/kg body weight; Sigma-Aldrich), colistin (4.2 mg/kg body weight; Sigma-Aldrich), metronidazole (21.5 mg/kg body weight; Sigma-Aldrich) and vancomycin (40 mg/kg body weight; Sigma-Aldrich) for 3 days by oral administration [31]. Two days after the antibiotic treatment, mice were intraperitoneally administrated with a single dose of clindamycin (10 mg/kg) and on the next day were infected orogastrically with 100 μL of PBS containing 5×10^7^ spores of *C. difficile* strain R20291. Mice were housed individually in sterile cages with *ad libitum* access to food and water. All procedures and mouse handling were performed aseptically in a biosafety cabinet to contain spore-mediated transmission.

The clinical condition of mice was monitored daily with a scoring system. The presence of diarrhea was classified according to severity as follows: (i) normal stool (score = 0); (ii) color change/consistency (score = 1); (iii) presence of wet tail or mucosa (score = 2); (iv) liquid stools (score = 3). A score higher than 1 was considered as diarrhea [49]. The animals were weighted daily after infection and other clinical symptoms as physical aspect (i.e., abnormal/hunched gait, piloerection), spontaneous behavior (i.e., lethargy, inactivity or lack of mobility) and emaciation were monitored as described [50]. Moribund mice or mice displaying overt signs of disease were sacrificed. At the time of sacrifice, ileum, proximal, median and distal colon were collected as well as the cecal content.

### Evaluation of BclA3_CTD_-specific IgG levels in mice serum

The blood collected the day before each immunization, two days before and at the time of sacrifice was incubated at 37°C for 30 minutes and then centrifuged at 5 000 rpm for 10 minutes at 4°C. The supernatant, containing the serum fraction was stored at −20°C until use. To assess the production of IgG against BclA3_CTD_, an Enzyme-linked immunosorbent assay (ELISA) was performed. Purified BclA3_CTD_ (50 ng/well), spores of *C. difficile* R20291 (1.6×10^7^ spores/well) or *C. difficile* R20291Δ*bcla3* (1.6×10^7^ spores/well) were coated onto 96-wells plates and incubated overnight at 4°C. Then, the samples were blocked with PBS-0.05% Tween-20 (PBS-T) containing 2% BSA for 1 hour at 37°C. After several washes, the wells were next incubated with 1:100 of animal serum (in 1% BSA in PBS-T). The plates were incubated 2 hours at 37°C. After the removal of non-adherent IgG by several washes, the plates were incubated with 1:5 000 secondary antibody anti-IgG mouse HRP, for 1 hour at 37°C. Finally, the colorimetric reaction was initiated upon the addition of 50 μL of reaction buffer containing 0.05 M citric acid, 0.1 M disodiumhydrogen phosphate, 2 mg/mL of o-phenlyendiamine (Sigma-Aldrich, U.S.A.) and 0.015% of H_2_O_2_ (Merck, Germany). The reaction was stopped after 20 minutes with 25 μL of 4,5 N of H_2_SO_4_ and absorbance was measured at 492 nm. The experiment was performed in duplicate.

### Quantification of *C. difficile* spores from feces and colon samples

Fecal samples were collected in the following five days after infection and were stored at −20°C until *C. difficile* spore quantification. On the day of the analysis, 10 μL of PBS was added for each mg of stools, mixed and incubated for 30 minutes at room temperature. Then, 50 μL of absolute ethanol (Sigma-Aldrich) was added to 50 μL of feces which were incubated for 30 minutes at room temperature.

Samples were serially diluted and plated onto selective medium supplemented with Taurocholate (0.1% w/v), Cefoxitin (16 μg/mL) and L-cycloserine (250 μg/mL) (TCCFA plates). The plates were incubated anaerobically at 37°C for 48 hours, the *C. difficile* colonies were counted, and the results were expressed as the Log_10_ of CFU/g of feces.

Proximal, median and distal colon were collected from mice upon sacrifice and washed with PBS with a syringe. They were posteriorly resuspended and homogenized with 2.5 µl of PBS for each mg of tissue. Upon incubation at room temperature with absolute ethanol and serially diluted they were plated onto TCCFA plates. The plates were incubated anaerobically at 37°C for 48 hours. Finally, the colony count was expressed as the Log_10_ of CFU/g of tissue.

### Cytotoxicity assay

Vero cell cytotoxicity was performed as described previously [51]. Briefly, 96-well flat-bottom microtiter plates were seeded with Vero cells at a density of 10^5^ cells/well. Mice caecum contents were kept at −20°C prior use. At the time of the experiment caecum contents were suspended in PBS (10 μL of PBS per mg of cecal content), vortexed and centrifuged (14 000 rpm, 5 minutes). Filter-sterilized supernatant was serially diluted in DMEM supplemented with 10% Fetal bovine serum (FBS) and 1% penicillium streptomycin. 100 μL of each dilution was added to wells containing Vero cells. Plates were screened for cell rounding 16 hours after incubation at 37°C. The cytotoxic titer was defined as the reciprocal of the highest dilution that produced rounding in at least 80% of Vero cells per gram of luminal samples under X200 magnification.

### Statistical analysis

Prism 8 (GraphPad Software, Inc.) was used for statistical analysis. Normality was assessed by Shapiro-Wilk test. For populations that did not follow a normal distribution significance between groups was assessed by Mann-Whitney unpaired t-test. Comparative analysis between groups was performed by analysis of variance with Turkey’s multiple comparison test for populations that followed a normal distribution. A P-value of ≤0.05 was accepted as the level of statistical significance.

## References

1. Kelly, C.P., C. Pothoulakis, and J.T. LaMont, Clostridioides difficile colitis. N Engl J Med, 1994. 330(4): p. 257–62.

2. Johnson, S., et al., Recurrences of Clostridioides difficile diarrhea not caused by the original infecting organism. J Infect Dis, 1989. 159(2): p. 340–3.

3. O’Neill, G.L., M.H. Beaman, and T.V. Riley, Relapse versus reinfection with Clostridioides difficile. Epidemiol Infect, 1991. 107(3): p. 627–35.

4. Wilcox, M.H., et al., Recurrence of symptoms in Clostridioides difficile infection--relapse or reinfection? J Hosp Infect, 1998. 38(2): p. 93–100.

5. Miller, B.A., et al., Comparison of the burdens of hospital-onset, healthcare facility-associated Clostridioides difficile Infection and of healthcare-associated infection due to methicillin-resistant Staphylococcus aureus in community hospitals. Infect Control Hosp Epidemiol, 2011. 32(4): p. 387–90.

6. Wiegand, P.N., et al., Clinical and economic burden of Clostridioides difficile infection in Europe: a systematic review of healthcare-facility-acquired infection. J Hosp Infect, 2012. 81(1): p. 1–14.

7. Zhang, S., et al., Cost of hospital management of Clostridioides difficile infection in United States-a meta-analysis and modelling study. BMC Infect Dis, 2016. 16(1): p. 447.

8. Escobar-Cortes, K., J. Barra-Carrasco, and D. Paredes-Sabja, Proteases and sonication specifically remove the exosporium layer of spores of Clostridioides difficile strain 630. J Microbiol Methods, 2013. 93(1): p. 25–31.

9. Joshi, L.T., et al., Contribution of spores to the ability of Clostridioides difficile to adhere to surfaces. Appl Environ Microbiol, 2012. 78(21): p. 7671–9.

10. Strong, P.C., et al., Identification and characterization of glycoproteins on the spore surface of Clostridioides difficile. J Bacteriol, 2014. 196(14): p. 2627–37.

11. Xue, Q., et al., Entry of Bacillus anthracis spores into epithelial cells is mediated by the spore surface protein BclA, integrin alpha2beta1 and complement component C1q. Cell Microbiol, 2011. 13(4): p. 620–34.

12. Barra-Carrasco, J., et al., The Clostridioides difficile exosporium cysteine (CdeC)-rich protein is required for exosporium morphogenesis and coat assembly. J Bacteriol, 2013. 195(17): p. 3863–75.

13. Pizarro-Guajardo, M., et al., Ultrastructural Variability of the Exosporium Layer of Clostridioides difficile Spores. Appl Environ Microbiol, 2016. 82(7): p. 2202–2209.

14. Mora-Uribe, P., et al., Characterization of the Adherence of Clostridioides difficile Spores: The Integrity of the Outermost Layer Affects Adherence Properties of Spores of the Epidemic Strain R20291 to Components of the Intestinal Mucosa. Front Cell Infect Microbiol, 2016. 6: p. 99.

15. Ghose, C., et al., Immunogenicity and protective efficacy of Clostridioides difficile spore proteins. Anaerobe, 2016. 37: p. 85–95.

16. Phetcharaburanin, J., et al., The spore-associated protein BclA1 affects the susceptibility of animals to colonization and infection by Clostridioides difficile. Mol Microbiol, 2014. 92(5): p. 1025–38.

17. Pizarro-Guajardo, M., et al., Characterization of the collagen-like exosporium protein, BclA1, of Clostridioides difficile spores. Anaerobe, 2014. 25: p. 18–30.

18. Maia, A.R., et al., Induction of a Specific Humoral Immune Response by Nasal Delivery of Bcla2ctd of Clostridioides difficile. Int J Mol Sci, 2020. 21(4).

19. Aubry, A., et al., In vitro Production and Immunogenicity of a Clostridioides Difficile Spore-Specific BclA3 Glycopeptide Conjugate Vaccine. Vaccines (Basel), 2020. 8(1).

20. Cutting, S.M., et al., Oral vaccine delivery by recombinant spore probiotics. Int Rev Immunol, 2009. 28(6): p. 487–505.

21. Huang, J.M., et al., Mucosal delivery of antigens using adsorption to bacterial spores. Vaccine, 2010. 28(4): p. 1021–30.

22. de Souza, R.D., et al., Bacillus subtilis spores as vaccine adjuvants: further insights into the mechanisms of action. PLoS One, 2014. 9(1): p. e87454.

23. Isticato, R., et al., Surface display of recombinant proteins on Bacillus subtilis spores. J Bacteriol, 2001. 183(21): p. 6294–301.

24. Hoang, T.H., et al., Recombinant Bacillus subtilis expressing the Clostridioides perfringens alpha toxoid is a candidate orally delivered vaccine against necrotic enteritis. Infect Immun, 2008. 76(11): p. 5257–65.

25. Hinc, K., et al., Expression and display of UreA of Helicobacter acinonychis on the surface of Bacillus subtilis spores. Microb Cell Fact, 2010. 9: p. 2.

26. Permpoonpattana, P., et al., Immunization with Bacillus spores expressing toxin A peptide repeats protects against infection with Clostridioides difficile strains producing toxins A and B. Infect Immun, 2011. 79(6): p. 2295–302.

27. Ning, D., et al., Surface-displayed VP28 on Bacillus subtilis spores induce protection against white spot syndrome virus in crayfish by oral administration. J Appl Microbiol, 2011. 111(6): p. 1327–36.

28. Youngman, P., J.B. Perkins, and R. Losick, Construction of a cloning site near one end of Tn917 into which foreign DNA may be inserted without affecting transposition in Bacillus subtilis or expression of the transposon-borne erm gene. Plasmid, 1984. 12(1): p. 1–9.

29. Naclerio, G., et al., Bacillus subtilis spore coat assembly requires cotH gene expression. J Bacteriol, 1996. 178(15): p. 4375–80.

30. Zilhao, R., et al., Assembly and function of a spore coat-associated transglutaminase of Bacillus subtilis. J Bacteriol, 2005. 187(22): p. 7753–64.

31. Chen, X., et al., A mouse model of Clostridioides difficile-associated disease. Gastroenterology, 2008. 135(6): p. 1984–92.

32. Pizarro-Guajardo, M., et al., New insights for vaccine development against Clostridioides difficile infections. Anaerobe, 2019. 58: p. 73–79.

33. Sorman, A., et al., How antibodies use complement to regulate antibody responses. Mol Immunol, 2014. 61(2): p. 79–88.

34. Yu, L.H. and S.M. Cutting, The effect of anti-spore antibody responses on the use of spores for vaccine delivery. Vaccine, 2009. 27(34): p. 4576–84.

35. Oggioni, M.R., et al., Bacillus spores for vaccine delivery. Vaccine, 2003. 21 Suppl 2: p. S96–101.

36. Duc le, H., et al., Bacterial spores as vaccine vehicles. Infect Immun, 2003. 71(5): p. 2810–8.

37. Rhee, K.J., et al., Role of commensal bacteria in development of gut-associated lymphoid tissues and preimmune antibody repertoire. J Immunol, 2004. 172(2): p. 1118–24.

38. Sibley, L., et al., Recombinant Bacillus subtilis spores expressing MPT64 evaluated as a vaccine against tuberculosis in the murine model. FEMS Microbiol Lett, 2014. 358(2): p. 170–9.

39. Wang, J., et al., Intranasal administration with recombinant Bacillus subtilis induces strong mucosal immune responses against pseudorabies. Microb Cell Fact, 2019. 18(1): p. 103.

40. Tavares Batista, M., et al., Gut adhesive Bacillus subtilis spores as a platform for mucosal delivery of antigens. Infect Immun, 2014. 82(4): p. 1414–23.

41. Isticato, R. and E. Ricca, Spore Surface Display. Microbiol Spectr, 2014. 2(5).

42. Duc le, H., et al., Intracellular fate and immunogenicity of B. subtilis spores. Vaccine, 2004. 22(15-16): p. 1873–85.

43. Casula, G. and S.M. Cutting, Bacillus probiotics: spore germination in the gastrointestinal tract. Appl Environ Microbiol, 2002. 68(5): p. 2344–52.

44. Hoa, T.T., et al., Fate and dissemination of Bacillus subtilis spores in a murine model. Appl Environ Microbiol, 2001. 67(9): p. 3819–23.

45. Harwood, C.R. and S.M. Cutting, Molecular biological methods for Bacillus. Modern microbiological methods. 1990, Chichester; New York: Wiley. xxxv, 581 p.

46. Calderon-Romero, P., et al., Clostridioides difficile exosporium cysteine-rich proteins are essential for the morphogenesis of the exosporium layer, spore resistance, and affect C. difficile pathogenesis. PLoS Pathog, 2018. 14(8): p. e1007199.

47. Edwards, A.N. and S.M. McBride, Isolating and Purifying Clostridioides difficile Spores. Methods Mol Biol, 2016. 1476: p. 117–28.

48. Isticato, R., E. Ricca, and L. Baccigalupi, Spore Adsorption as a Nonrecombinant Display System for Enzymes and Antigens. J Vis Exp, 2019(145).

49. Warren, C.A., et al., Vancomycin treatment’s association with delayed intestinal tissue injury, clostridial overgrowth, and recurrence of Clostridioides difficile infection in mice. Antimicrob Agents Chemother, 2013. 57(2): p. 689–96.

50. Deakin, L.J., et al., The Clostridioides difficile spo0A gene is a persistence and transmission factor. Infect Immun, 2012. 80(8): p. 2704–11.

51. Theriot, C.M., et al., Cefoperazone-treated mice as an experimental platform to assess differential virulence of Clostridioides difficile strains. Gut Microbes, 2011. 2(6): p. 326–34.

